# Lipid digestion and autophagy are critical energy providers during acute glucose depletion in *Saccharomyces cerevisiae*

**DOI:** 10.1101/728667

**Authors:** Carmen A. Weber, Karthik Sekar, Jeffrey H. Tang, Philipp Warmer, Uwe Sauer, Karsten Weis

## Abstract

The ability to tolerate and thrive in diverse environments is paramount to all living organisms, and many organisms spend a large part of their lifetime in starvation. Upon acute glucose starvation, yeast cells undergo drastic physiological and metabolic changes and reestablish a constant - though lower – level of energy production within minutes. The molecules that are rapidly metabolized to fuel energy production under these conditions are unknown. Here, we combine metabolomics and genetics, to characterize the cells’ response to acute glucose depletion and identify pathways that ensure survival during starvation. We show that the ability to respire is essential for maintaining the energy status and to ensure viability during starvation. Measuring the cells’ immediate metabolic response, we find that central metabolites drastically deplete and that the intracellular AMP to ATP ratio strongly increases within 20-30 seconds. Furthermore, we detect changes in both amino acid and lipid metabolite levels. Consistent with this, bulk autophagy, a process that frees amino acids, as well as lipid degradation via β-oxidation contribute in parallel to energy maintenance upon acute starvation. In addition, both these pathways ensure long-term survival during starvation. Thus, our results identify bulk autophagy and β-oxidation as important energy providers during acute glucose starvation.

## Introduction

Cellular life depends on metabolic substrates for growth and survival. Glucose is a common substrate and convenient energy source that feeds directly into glycolysis, leading to rapid cell growth and proliferation in many microbes. Budding yeast, for example, depend highly on glucose as an energy source for rapid growth. One of their evolutionary advantages is that they can outgrow many of their competitors when supplied with glucose. The dependence on glucose and the ability to respond to glucose starvation is not only common in microbes and unicellular organisms, but also pertains to multicellular eukaryotes. A large number of cancer cells for instance rely mostly on glucose and glycolysis even when oxygen is present (i.e. the Warburg effect), which leads to outcompeting of non-cancerous cells in terms of proliferation. Not only cancerous cells rely predominantly on glucose, but also healthy cell types such as neurons, which depend exclusively on glucose as their energy source. Interestingly, energy deficits and low glucose levels have been correlated with neurodegenerative diseases such as Alzheimer’s (1–3). Thus, understanding how cells cope with starvation is crucial for elucidating both normal cellular processes as well as aberrant behaviors in disease.

In microbes such as the budding yeast *Saccharomyces cerevisiae*, starvation is especially ubiquitous (4), and must be counteracted very rapidly. Budding yeast have developed an intricate and rapid response to glucose starvation, including reduction of transcriptional and translational activity (5–7), autophagy of cytoplasmic components and lipids for energetic needs (8), and reduction of macromolecular diffusivity (9, 10). Starvation states can differ depending on the type of starvation or nutrient limitation (11).

Previous studies have predominantly focused on starvation-induced metabolic changes that occur over hours or longer (8, 12, 13). Yet, metabolic pools rapidly deplete and fill within seconds (14, 15) and any initial responses have likely been missed in these studies. Furthermore, metabolic changes can occur much faster than transcriptional and protein abundance changes (16), and metabolism is therefore likely amongst the first responders to changes in the environment. However, the rapid metabolic changes that occur at the onset of starvation have remained unclear, and the critical energy sources that are utilized upon acute glucose starvation – leading to a new energy equilibrium within minutes (9) - have not been well characterized. Additionally, it is poorly understood how metabolism communicates with the rest of the cell to ensure long-term survival.

To begin to address some of these questions, we performed a metabolomic analysis upon acute glucose starvation in budding yeast. Our results reveal that upper glycolysis metabolites deplete within seconds of acute glucose starvation while intracellular lipid pools increase or are maintained. In addition, we found that the metabolic flow through amino acid metabolites changes significantly. Based on these findings we assessed the effect of lipid digestion and autophagy on the cellular energy status upon acute glucose starvation and found that lipid degradation and autophagy act together to ensure ATP maintenance as well as cellular survival in acute glucose starvation.

## Results

### Respiration is crucial for survival and energy maintenance upon acute glucose starvation

Our previous results suggested that respiration is critical for energy production upon acute glucose withdrawal (9). To address this question of how important respiration activity is for the acute glucose starvation response we genetically deleted *CBP2* in order to abolish aerobic respiration. The *CBP2* gene product facilitates the splicing of the cytochrome B oxidase pre-RNA (17). Therefore, a knockout of *CBP2* inhibits the ability of cells to respire. In *cbp2Δ* mutants, intracellular ATP levels dropped to non-detectable levels within 1 h of acute glucose starvation and remained immeasurable until 19 h. Furthermore, the survival rate in the absence of glucose was dramatically reduced compared to the wild-type (Figure 1a). This showed that respiratory activity is critical to provide the necessary energy to promote survival during starvation.

**Figure 1.**
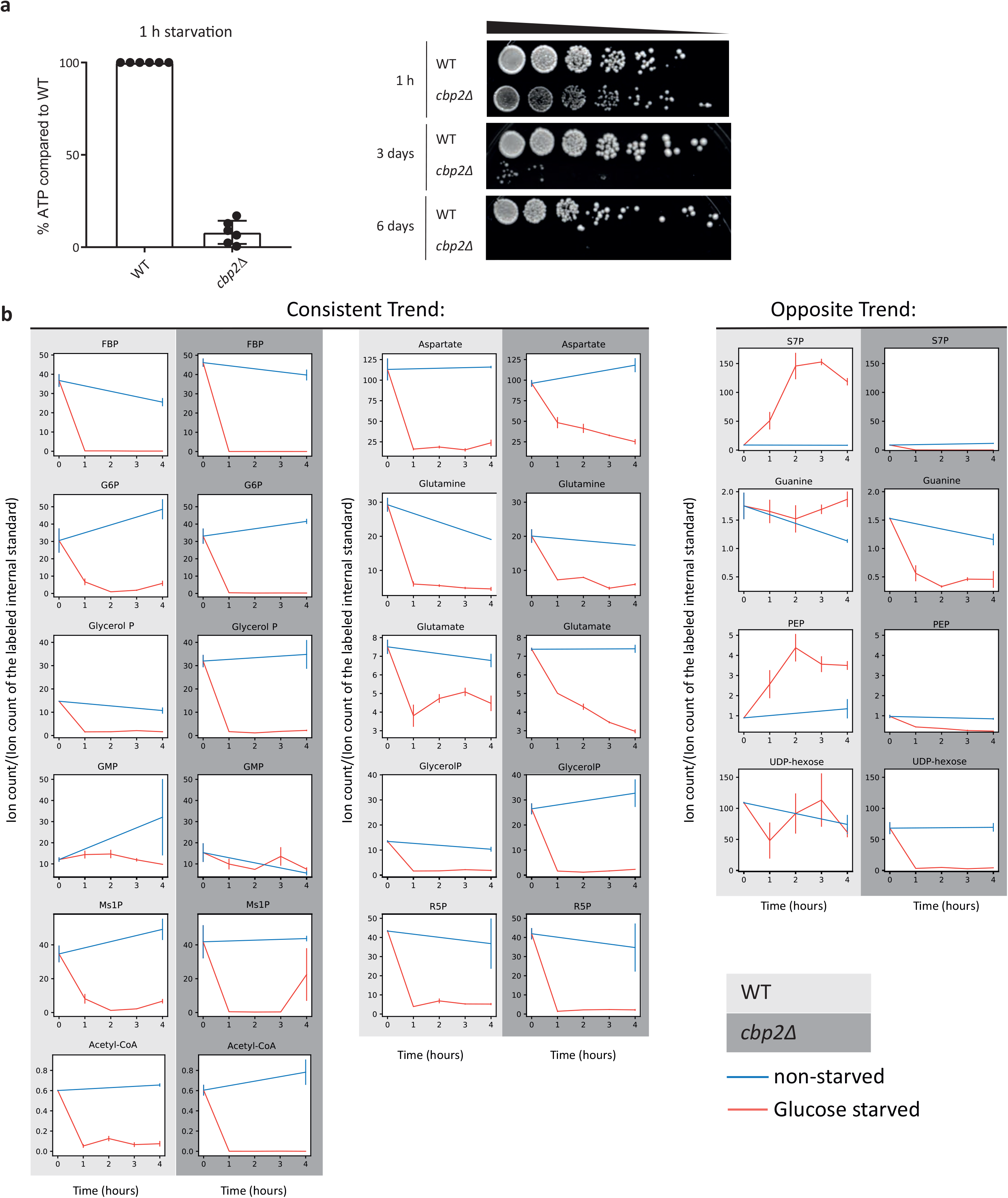
The ability to respire is crucial for survival and energy maintenance upon acute glucose starvation. **(a)** Relative ATP levels in the respiratory deficient *cbp2Δ* mutant compared to WT after 1 h of acute glucose starvation. Mean, standard deviation, and biological replicates are shown. Right panel: Survival after 1h, 3 days and 6 days of acute starvation **(b)** Change in ion intensity for metabolites after 0, 1, 2, 3, and 4 h of acute glucose starvation (red, glucose starved), compared to non-starved cells (blue, non-starved). Comparison between wild-type (WT) cells and *cbp2Δ* cells. Abbreviations: FBP - fructose bisphosphate, G6P – Glucose-6-phosphate, Glycerol P – Glycerol phosphate, GMP - guanosine monophosphate, Ms1P -mannose 1-phosphate, R5P – Ribose-5-phosphate, S7P - sedoheptulose-7-guanine, PEP – phosphoenolpyruvate. Average and standard error (error bar) of 2 biological replicates are shown.

### Metabolite pools deplete globally within 1 h of starvation in cells unable to respire

Having established that respiration is required for energy maintenance and survival upon glucose starvation (Figure 1a), we next aimed to identify the substrate(s) feeding respiration in these conditions. We hypothesized that respiration substrates cannot be efficiently metabolized in a respiration-deficient mutant and hence might accumulate in these cells. We therefore measured and compared central carbon metabolites between wild-type and *cbp2Δ* cells on a time scale of 1-4 h of starvation (Figure 1b, Figure S1). A large swath of intracellular metabolites exhibited similar depletion patterns between the two strains, including many central metabolites in glycolysis (e.g. glycerol-phosphate, fructose-bisphosphate, and glucose-6-phosphate) and in the citric acid cycle (e.g. acetyl-CoA, glutamate, glutamine). Some select metabolites, such as glucose-6-phosphate, acetyl-CoA, ribose-5-phosphate, and glutamate, depleted more completely in the *cbp2Δ* mutants compared to wild-type, and others, including sedoheptulose-7-guanine, phosphoenolpyruvate, and UDP-hexose, diminished entirely during glucose starvation in *cbp2Δ* but remained constant or even accumulated in wild-type cells. However, we could not detect any intracellular metabolite that specifically accumulated in *cbp2Δ* cells upon glucose starvation. We conclude that intracellular metabolite pools deplete rapidly in both wild-type and respiration deficient cells, and do so even more rapidly or more completely if cells are unable to respire.

### Maintenance of ATP levels and increased survival under acute glucose starvation are neither explained by extracellular amino acids nor intracellular glycogen

Our initial analyses of intracellular metabolites did not provide us with obvious candidates fueling respiration that became apparent through accumulation in a respiration deficient mutant. We next focused our attention on candidate substrates during starvation, namely extracellular metabolites as well as intracellular glycogen reserves.

Extracellular metabolites could either be used as an external energy source or point to the activity of intracellular pathways based on secreted metabolites. To examine changes in extracellular metabolites, we grew cells to OD 0.5-0.8 in synthetic complete media with glucose (SCD), essential amino acids, and a yeast nitrogen source (see Materials and Methods for details). The cells were then acutely starved by washing into the same synthetic media lacking glucose. The supernatant was sampled before and after the transition, and a control experiment was conducted by washing the cells with SCD to ensure that medium change itself did not lead to secondary metabolic effects. The strongest effect we observed was a rapid depletion of extracellular amino acids within the first hour of acute glucose starvation, specifically, two exemplary amino acids, aspartate and methionine (Figure 2a). Monitoring their abundances beyond this first hour, no additional appreciable changes could be observed for up to 19 h. Since our glucose starvation medium contains essential amino acids, this could suggest that extracellular amino acids are assimilated and catabolized during acute starvation. To test whether the extracellular amino acids serve as energy sources on a short time scale during glucose starvation, we measured intracellular ATP levels within the uptake period of 1 h or within longer starvation up to 19 h in cells starved from glucose (SC), starved from glucose and amino acids (SC −AA), and cells starved in water (H_2_O) (Figure 2c). The ATP levels did not drop much lower in water nor in the medium lacking glucose and amino acids compared to medium only lacking glucose, neither for rapid (30 min-1 h) nor short (19 h) starvation duration.

**Figure 2.**
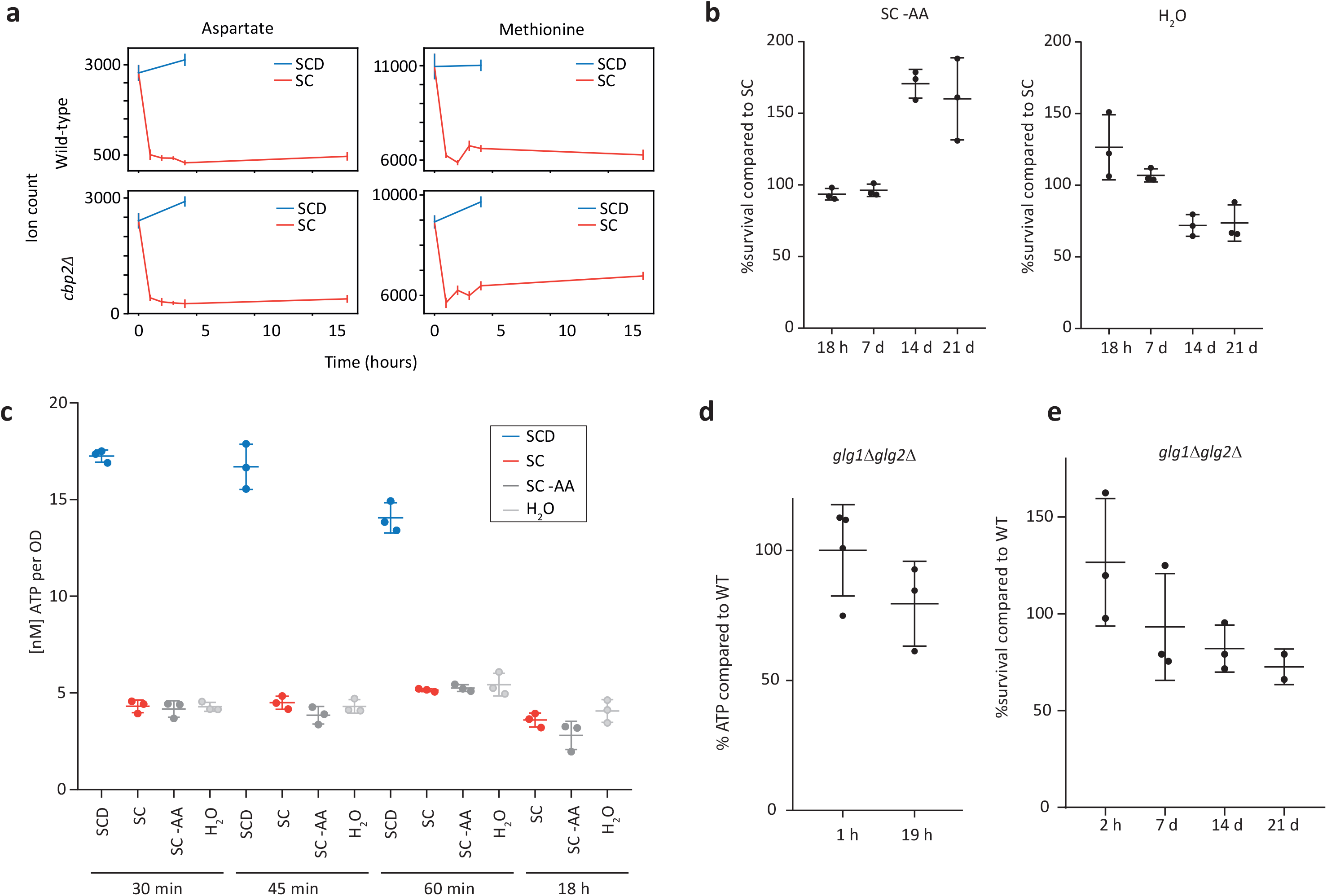
Extracellular aminoacids and intracellular glycogen levels are not the main short-term energy source upon acute glucose starvation. **(a)** Extracellular aspartate and methionine primarily deplete within the first 1 h of starvation. Wild-type and *cbp2Δ* mutants were grown in SCD (synthetic complete medium with glucose) and then switched to SCD or SC (synthetic complete medium without glucose). Extracellular samples were taken and measured for relative abundance of ions corresponding to aspartate and methionine. Error bars indicate the standard error of two biological replicates. **(b)** Survival of wild-type yeast cells after 18 h - 21 days of acute glucose and amino acid starvation (SC -AA) and acute starvation in water (H_2_O), compared to survival of cells only starved from glucose (SC) **(c)** ATP levels after hours and minutes of starvation in SC, SC-AA, or H_2_O. **(d)** ATP levels of *glg1Δglg2Δ* mutants compared to WT after 19 h of acute glucose starvation. **(e)** Quantified spotting assays after 2h-21 days of starvation. Mean and standard deviation and values of biological replicates are shown.

To complement the ATP measurements with functional evidence, we also conducted spotting assays for cells starved in SC media, in SC media without amino acids (SC-AA), and in water (H_2_O) to assess survival and recovery after the starvation stress. Cells starved from both glucose and amino acids initially showed no survival difference compared to cells starved from glucose only (up to day 7 of starvation) and even seemed to have a survival advantage after 14 days of starvation (Figure 2b, Figure S2). Survival of cells washed into water instead of SC or SC-AA was only slightly and insignificantly impaired after 14-21 days compared to the glucose-starved cells (Figure 2b, Figure S2). Although extracellular amino acids were taken up within 1 h of starvation, we thus conclude that they did neither affect ATP levels upon acute glucose starvation nor provide cells with a survival advantage long-term.

Another potential candidate substrate during glucose starvation is intracellular glycogen. In various starvation conditions, intracellular glycogen resources were proposed to build up or to be used for energy maintenance (18). Since glycogen was shown to decrease to non-measurable levels after glucose starvation (19), we tested whether intracellular glycogen resources sustain ATP maintenance and survival during short-term starvation. However, a mutant deficient in glycogen synthesis (*glg1Δglg2Δ*) did not adversely affect ATP levels after 1-19 h of acute glucose starvation, and showed no significant survival deficiency even after 21 days (Figure 2d,e, Figure S4). Thus, glycogen stores in the cell are not a main energy source upon acute glucose starvation.

### Central carbon metabolites exhibit sub-minute changes to glucose deprivation

Our initial survey did not provide us with good candidate substrates that yeast cells utilize to ensure their survival upon acute glucose starvation, but suggests that many major metabolic changes occur on a timescale faster than 1 hour. Hence, our experiments thus far might have missed initial responses that could be important for long term survival. Therefore, we expanded our metabolomic analyses to earlier time points. Since metabolic pools can deplete and fill within seconds (14, 15), we aimed for measurements on a 10-60 second timescale. We employed a high-throughput, untargeted mass spectrometry approach (15, 20) to measure the global, starvation-induced metabolic changes that occur in budding yeast. Yeast cells were cultivated in SCD media, and using a rapid fast filtration setup (21, 22), we dispensed approximately 1 OD unit of yeast on a filter, perfused with SCD media for at least 10 s followed by exposure to media without glucose (SC) for varying times (10, 20, 30, and 60 s) (Figure 3a). The yeast containing filters were rapidly quenched after the designated exposure time, and intracellular metabolites were extracted. Relative changes of intracellular metabolite concentrations were measured and annotated (20). This semi-quantitative method allowed us to measure the response of approximately 350 ions that can be associated to 650 metabolites known to the metabolome of *S. cerevisiae* (23). Rapid, fold changes occurred in many metabolites visualized on a metabolic map (Figure 3b). All measured ion data is available in Supplementary Data 1.

**Figure 3:**
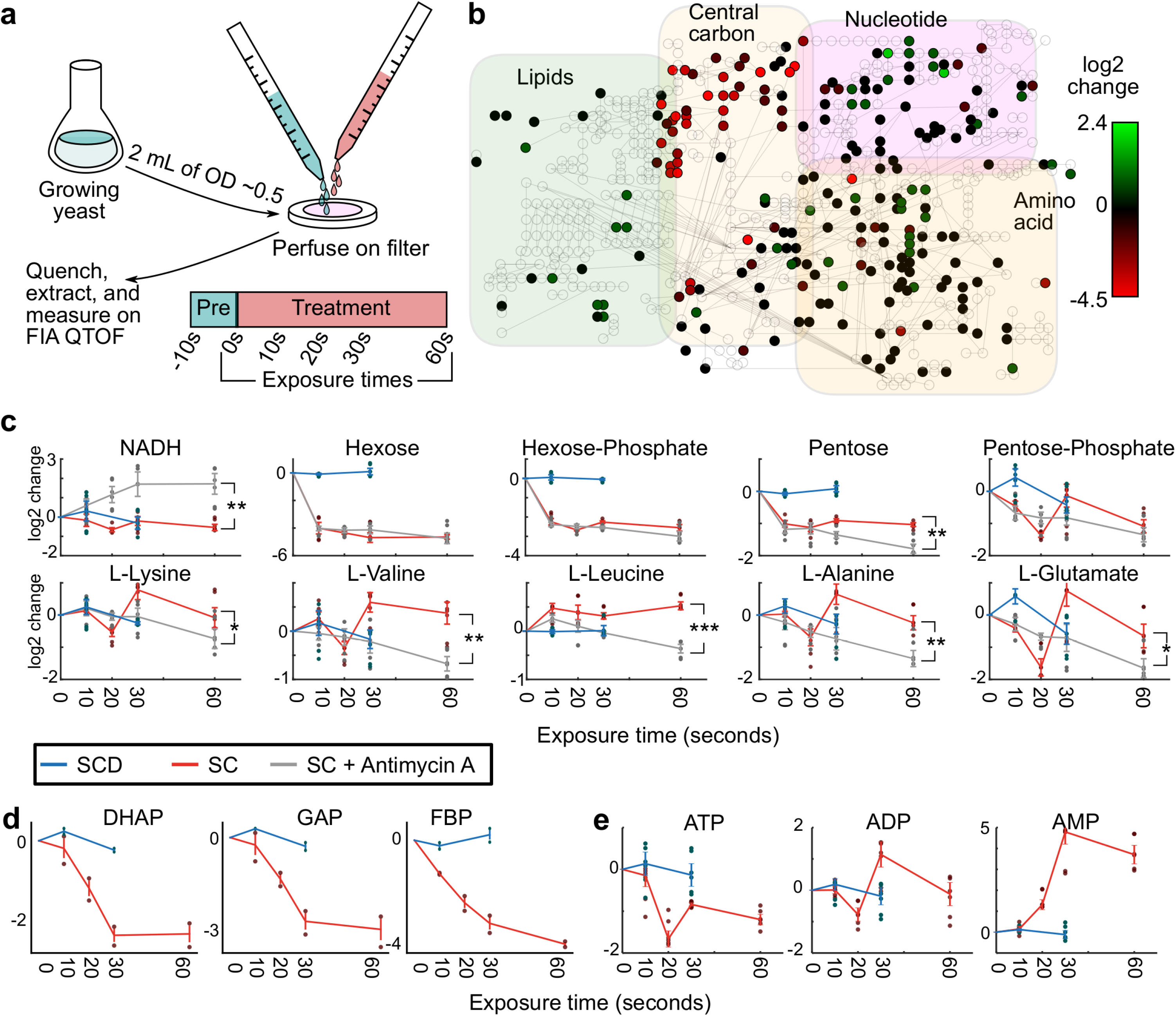
Central carbon metabolites exhibit sub-minute changes to glucose deprivation. **(a)** Fast filtration setup. Growing yeast (approximately 1 OD*mL) were deposited onto filters and quickly perfused with pretreatment media (SCD). At the 0 second timepoint, cells were perfused with either SCD or SC (starvation) medium, and after exposure for a given amount of time (10 s, 20 s, 30 s, and 60 s), the filters were rapidly quenched in extraction solution and measured on mass spectrometry for small metabolites (m/z <1000 g/mol). **(b)** A metabolic map of central carbon metabolism shows rapid depletion of metabolites in upper glycolysis and increase in lipid synthesis. The log2 change in ion intensity is shown between 60 s exposure of SC media versus the average of the 10 s and 30 s exposure on SCD media. **(c)** Glycolytic metabolites deplete strongly within 60 s upon exposure to SC media. The log2 adjusted values of intensity for specific ions are shown and labeled according to the annotated compound. Measurement time indicates the exposure time for the given media for the cells on the filter (SCD - blue; SC −red; SC with Antimycin A - gray). 6 dots are shown for each timepoint (3 biological replicates and 2 technical measurement replicates). Timepoint 0 s is extrapolated from the average of the SCD condition. Error bars indicate the standard error for three biological replicates. Significances are shown between the SC and SC with Antimycin A condition for time 60 s (* indicates 0.01 < *P* < 0.05, ** for 10^−4^ < *P* < 0.01, and *** for P < 10^−4^). **(d)** Measurement of other metabolites (dihydroxyacetone phosphate - DHAP, glyceraldehyde 3-phosphate - GAP, and fructose bisphosphate - FBP) with another mass spectrometer quantification method also corroborate the depletion of upper glycolysis metabolites. 2 dots are shown for each timepoint (2 biological replicates). Error bars indicate the standard error for two biological replicates. **(e)** ATP, ADP, and AMP levels as indicators of cellular energy homeostasis rapidly change during starvation.

How does respiration affect the metabolism at these early time points? To answer this question, we treated cells with Antimycin A and measured them in parallel with untreated cells in our fast filtration setup. Antimycin A was chosen as it specifically and rapidly inhibits the electron transport chain in mitochondria avoiding any potential adaptation effects that might have occurred in the long-term absence of respiration in *cbp2Δ* cells. An increase of NADH in Antimycin A treated cells confirmed the successful application of the drug, since the block of the electron transport chain inhibits NADH oxidation (25) (Figure 3c). Glucose, as indicated by the hexose ion, decreased within the first 10 seconds in both starvation conditions, indicating that the cells use up internal glucose very rapidly upon starvation.

Furthermore, the Antimycin A treatment in starvation led to a stronger decrease of most amino acid pools compared to cells that were only starved (Figure 3c, Figure S3), indicating that these metabolite pools are either degraded and potentially explored for energy generation much faster than in cells capable of respiration, or that protein degradation slows down in Antimycin A treated cells, leading to less generation of amino acids. These observations echoed our earlier observations about respiratory-deficient cells with generally lower metabolic pools over long time scales.

Glycolysis metabolites such as hexose phosphates decreased in both conditions within 10 seconds. Intermediates of the pentose phosphate pathway such as ribose phosphates (corresponding to the pentose phosphate ion) and potential precursors such as D-ribose (pentose) also decreased in both conditions, but were then maintained at lower levels in the untreated cells while they depleted even more in the respiratory deficient condition (Figure 3c) correlating with the increased phosphoenolpyruvate levels observed in the *cbp2Δ* cells at later timepoints (Figure 1b).

To further substantiate these observations in wild-type untreated cells, we obtained absolute metabolite concentrations with a targeted mass-spectrometry method (24) and found other glycolytic and pentose phosphate pathway metabolites to exhibit a similar rapid depletion (Figure 3d), specifically dihydroxyacetone phosphate, glyceraldehyde 3-phosphate, and fructose bisphosphate. The rapid depletion was comparable to an earlier study, examining *E. coli* carbon starvation (14). These results suggested that upon removal of the glucose input downstream metabolic activity continues, leading to the successive depletion of metabolic pools. Such depletion is expected to start with upper glycolysis metabolites as the entry point for glucose.

The energy charges of the cells (e.g. ATP, ADP, and AMP) showed similar rapid changes (Figure 3e). ATP depleted within seconds, and AMP increased in a near equivalent time frame. Our data suggest that global metabolic changes occur within seconds of starvation. Interestingly, we observed that both intracellular amino acid as well as lipid pools changed considerably within these initial time points. We therefore focused on these two classes for our further studies.

### Autophagy is important for energy maintenance upon acute glucose starvation and ensures long-term survival

Since our map of starvation-induced metabolite changes revealed changes of amino acid metabolites, we examined whether amino acid digestion could provide a pathway for ATP production through respiration, and whether intracellular proteins, which were synthesized prior to starvation, could be an amino acid source upon acute glucose removal. Since proteasomal degradation is one of the main specific protein degradation pathways, we explored whether proteasomal degradation contributed to ATP generation and maintenance within 1 h of glucose starvation. To test this, proteasomal activity was blocked using the small molecular inhibitor MG132. ATP concentrations in MG132-treated cells compared with the control cells showed no significant changes upon glucose starvation for 1 h (Figure 4a). Due to its potential cellular toxicity, we did not test the effect of MG132 on long-term survival, but our results nevertheless suggest that proteasomal degradation is not a significant energy source during the first hours of acute starvation.

**Figure 4.**
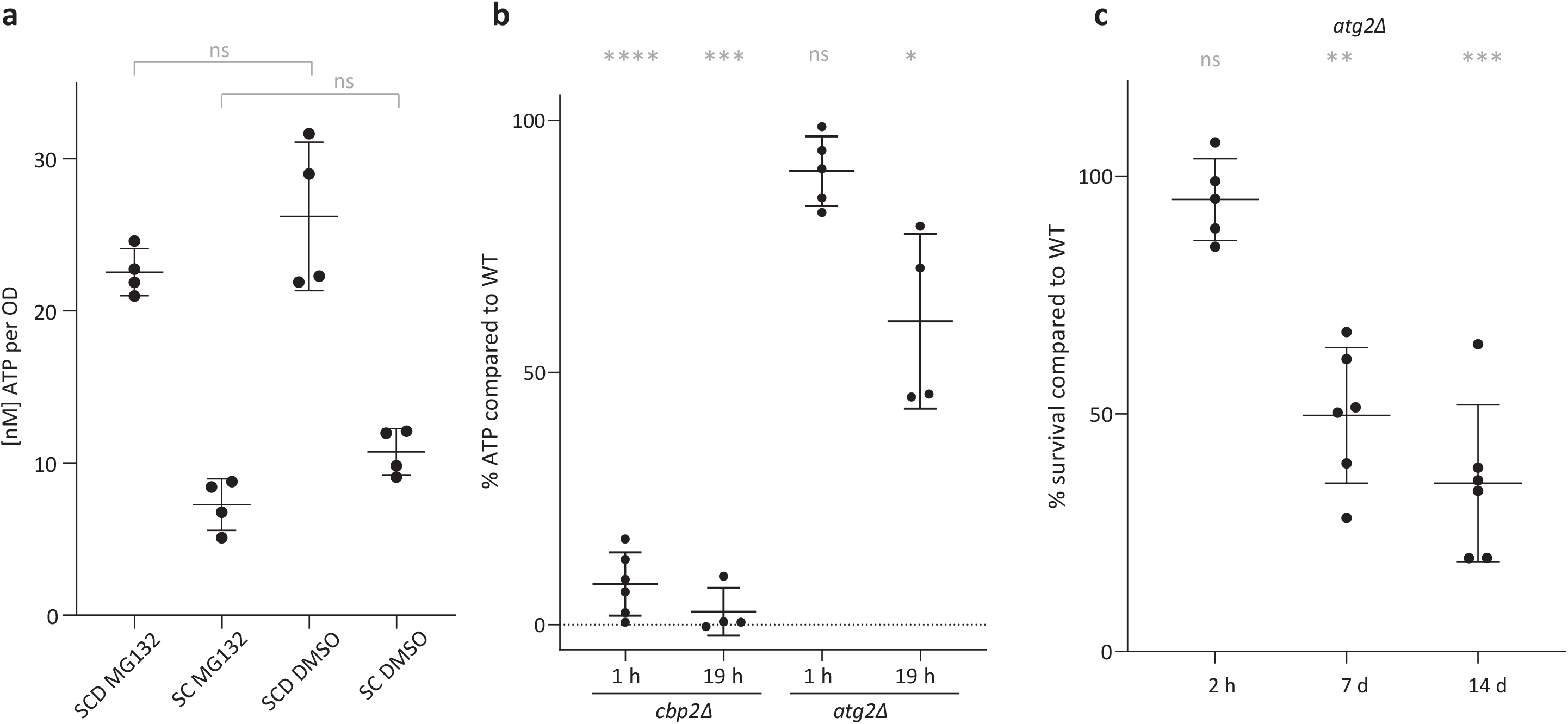
Unlike proteasomal activity, bulk autophagy contributes to survival and energy maintenance upon acute glucose starvation. **(a)** ATP levels after 1 h of pre-treatment with MG132 or DMSO as a control, followed by 1 h in acute SC or SCD medium with MG132/DMSO. **(b)** ATP levels in *cbp2Δ* and *atg2Δ* relative to wild-type (WT) cells after 1 h and 19 h of acute glucose starvation. **(c)** Survival of *atg2Δ* cells after 2 h to 14 days of glucose starvation. Means, standard deviations and biological replicate values are shown. Significances are shown between the WT and mutants (ns indicates non-significant, * indicates 0.05 > P > 0.01, ** indicates 0.01 > P > 0.001, *** indicates 0.001 > P > 0.0001, and **** indicates P < 0.0001). Significance was calculated using a one-way ANOVA test followed by a Holm-Sidak test.

Another pathway that frees building blocks and carbon substrates is bulk autophagy, which removes cytosolic components non-specifically triggering their degradation in the vacuole. We therefore examined whether bulk autophagy is important for energy maintenance. We generated a mutant incapable of organizing its pre-autophagosomal structure, *atg2Δ* (31). Since bulk autophagy and the cytoplasm-to-vacuole transport (CvT) pathway largely rely on the same machinery, including *ATG2*, the *atg2Δ* mutant will affect additionally any specific process that relies on the CvT for shuttling to the vacuole and degradation therein. ATP levels after 19 h of glucose starvation were significantly decreased in the *atg2Δ* mutant compared to WT cells (Figure 4b). In addition, fewer *atg2Δ* cells survived 7 days of glucose starvation in comparison to the isogenic wild-type control (Figure 4c, Figure S4). Thus, bulk autophagy as well as specific autophagic processes involving the CvT pathway contributed to energy maintenance and survival in acute glucose starvation.

Our data further implied that there must be alternative pathways to generate ATP in the short term as autophagy deficiency did not decrease ATP levels to the same levels as seen in the complete absence of respiration (Figure 4b) and did also not fully impair long term survival.

### β-oxidation is required for energy maintenance and survival upon acute glucose starvation together with autophagy

In addition to amino acids, lipid metabolite levels changed on these very rapid time scales and we observed a rapid increase in some lipids in wild-type untreated cells, such as hexadecanoic acid and 3-oxooctanoyl-CoA (Figure 5a). There are two potential non-exclusive models that could explain the changes we see. On one hand, since a majority of lipids in the cell originates from membranes, the observed changes could be caused by global membrane remodeling during starvation. On the other hand, the cells might liberate lipids to use them as an energy/carbon source during starvation, leading to an increase of free fatty acyls. To test the first model, we utilized a lipid extraction method designed for yeast (26) and measured the lipidome over time (Figure 5bc) sampling cells before glucose withdrawal and every 10 min after. We could annotate over 10,000 ions by exact mass matching to over 400 classes in the 8 categories within the LIPIDS MAPS scheme (27) and classified the different classes as increasing or decreasing (Figure 5b). In our dataset, the three main membrane lipid classes (sphingolipids, phospholipids and sterol lipids) were enriched consistent with the fact that the majority of lipids in cells originates from membranes (28). No major differences appeared between the lipids measured in control versus glucose starvation conditions. Nonetheless, a few traces were found to change and particularly, lipids annotated to dolichols and linear polyketides exhibited an upward trend during starvation (Figure 5d). While this suggested specific lipids and lipid classes may be synthesized or depleted in direct response to starvation, the overall lipid composition and therefore cellular membranes did not seem to appreciably change within rapid time scales (30 min or less). This argued that yeast cells do not undergo a major membrane remodeling during acute glucose starvation.

**Figure 5.**
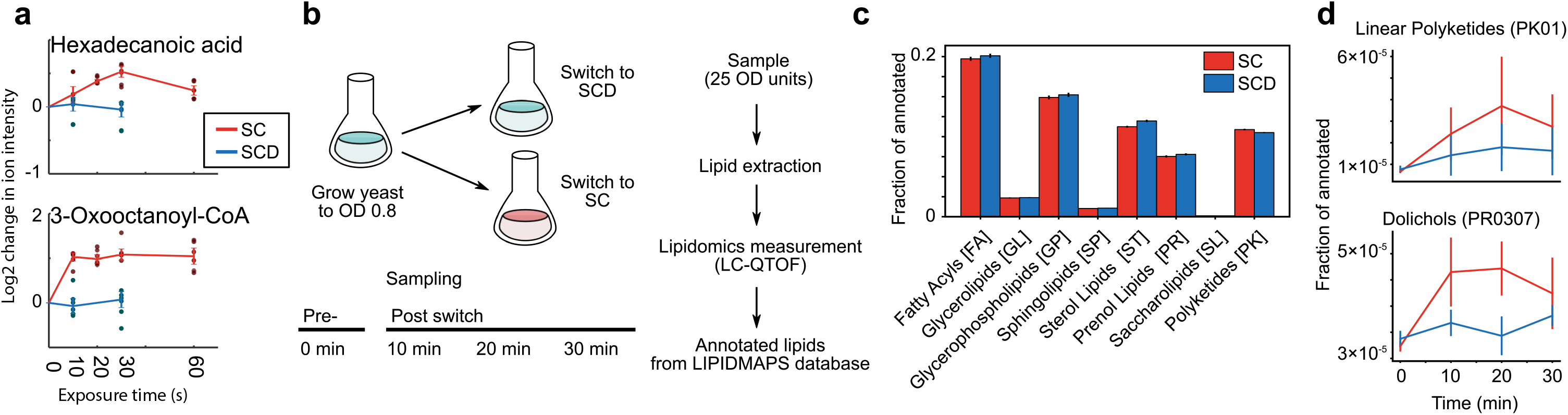
The lipidome during rapid starvation. **(a)** Hexadecanoid acid and 3-Oxooctanoyl-CoA as examples of lipid-related metabolites that accumulate within 10 to 60 seconds. The log2 adjusted values of intensity for specific ions are shown and labeled according to the annotated compound. Measurement time indicates the exposure time for the given media for the cells on the filter (SCD - blue; SC –red). 6 dots are shown for each timepoint (3 biological replicates and 2 technical measurement replicates). Timepoint 0 s is extrapolated from the average of the SCD condition. Error bars indicate the standard error for three biological replicates. **(b)** Experimental set up for measuring yeast lipidomics upon starvation entry. Exponentially growing yeast cells (OD 0.8 in SCD media), were sampled (time point 0 min), and then resuspended into fresh SCD or SC media. Samples were taken every 10 minutes thereafter. All samples were processed (see Methods) and measured. Putative lipids were annotated based on m/z and correspondence to the LIPIDMAPS database. **(c)** The normalized distribution of annotated lipids using the LIPIDMAPS identifiers. The major 8 lipid categories are shown. Error bars indicate the standard error between two biological replicates. **(d)** The lipid classes linear polyketides (PK01) and dolichols (PR0307) were identified as accumulating in glucose starvation compared to nutrient rich conditions. Error bars indicate the standard error between two biological replicates.

Since our results did not support the hypothesis that the changes we saw in lipid metabolites were caused by a major reorganization of cellular membranes, we hypothesized and examined whether the liberated lipids were used as an energy source to fuel respiration during starvation. μ-lipophagy was recently shown to play a role during long-term glucose restriction from 2% (w/v) to 0.4% (w/v) glucose (8). The study suggests that lipid droplets are digested through micro-lipophagy mediated by Atg14 to ensure survival in limiting glucose conditions. We therefore tested whether μ-lipophagy is also necessary for ATP level maintenance upon acute complete glucose starvation. Within 19 h of starvation, the ATP levels in *atg14Δ* mutants only deviated slightly from values measured in wild-type cells (Figure 6a). However, consistent with the results obtained by Seo et al., acute complete glucose starvation lead to survival deficiency after 7 days (Figure 6b). We conclude that μ - lipophagy by ATG14 did not contribute significantly to energy maintenance within the first 19 h of acute glucose starvation, but had long term effects on survival of the cells after 7 days.

**Figure 6.**
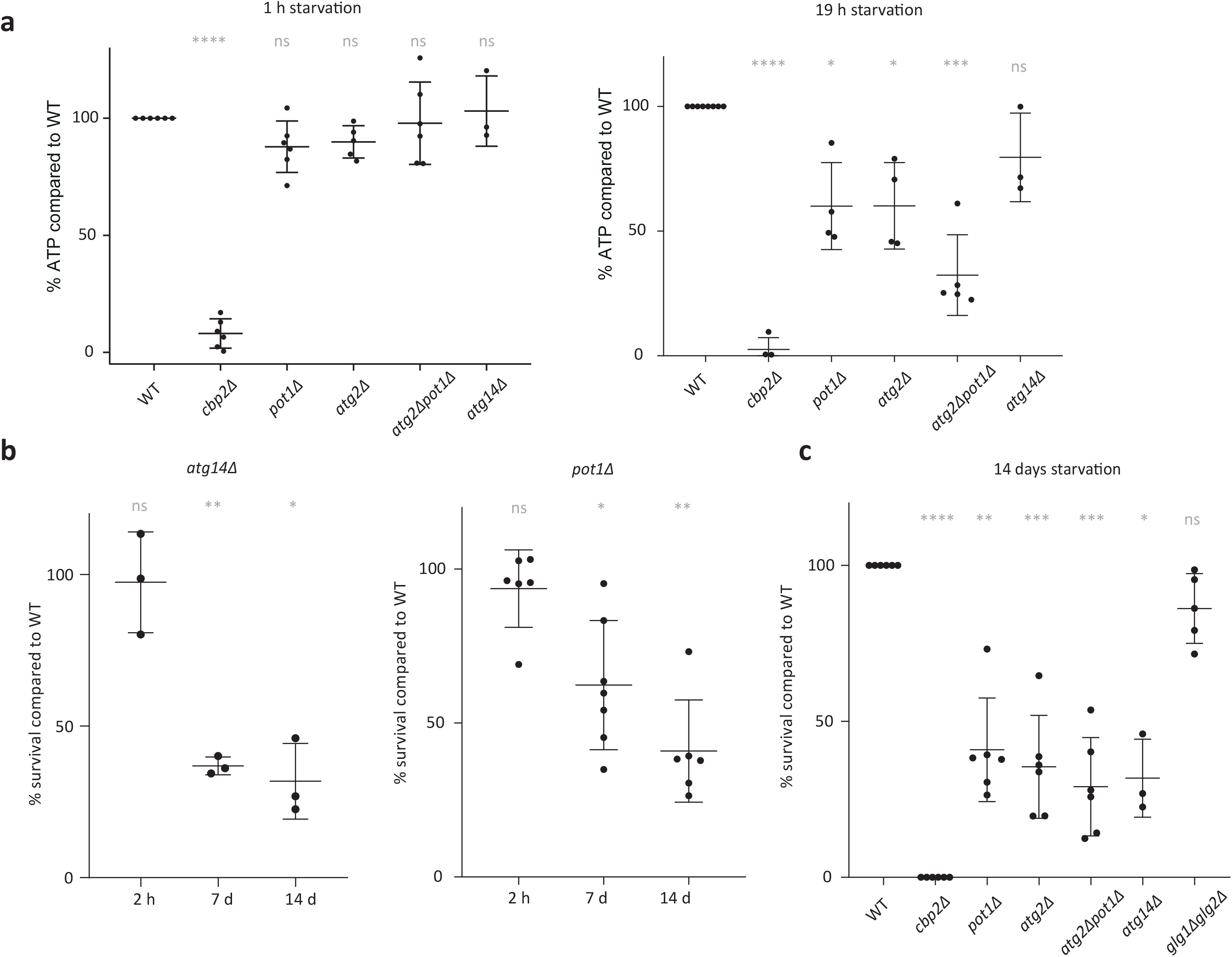
Lipid degradation and autophagy ensure survival and energy maintenance upon acute glucose starvation. **(a)** ATP levels in mutants compared to wild-type cells (WT) after 1 h or 19 h of acute glucose starvation. **(b)** Survival of *atg14Δ* and *pot1Δ* cells after 2 h - 14 days of acute glucose starvation. **(c)** Survival of mutant cells compared to wild-type (WT) cells after 14 days of acute glucose starvation. Significances are shown between the WT and mutants (ns indicates non-significant, * indicates 0.05 > P > 0.01, ** indicates 0.01 > P > 0.001, *** indicates 0.001 > P > 0.0001, and **** indicates P < 0.0001). Significance was calculated using a one-way ANOVA test followed by a Holm-Sidak test.

While μ-lipophagy specifically may only contribute to energy maintenance after the first day of starvation, general lipid digestion in peroxisomes could play a role early on. β-oxidation occurs exclusively in peroxisomes in yeast (29). To attenuate global lipid degradation in peroxisomes, we deleted the gene coding for Pot1. Pot1 is the only 3-ketoacyl-CoA thiolase in yeast that catalyzes the last step of β-oxidation, producing acetyl-CoA to feed into the citric acid cycle (30). Indeed, *pot1Δ* cells showed decreased ATP levels within 19 h of glucose starvation, demonstrating that β-oxidation of fatty acids contributes to ATP maintenance upon glucose starvation (Figure 6a). Furthermore, survival of *pot1Δ* cells decreased after 7-14 days in the absence of glucose (Figure 6bc, Figure S4).

Together our data suggest that lipid consumption by β-oxidation contributes to intracellular ATP levels upon acute glucose starvation within several hours of stress, and leads to a benefit for long term survival. By contrast, μ-lipophagy as a more specific way to consume lipids does not seem to contribute to short-term ATP maintenance, but is important for long term survival.

Similar to the results for the autophagy deficient mutant, the ATP and survival levels were not completely abolished in the *pot1Δ* mutant and did not reach the levels of the respiratory deficient *cbp2Δ* mutant. We therefore wondered whether bulk autophagy and β-oxidation might complement each other and generated a *pot1Δatg2Δ* double deletion strain. Consistent with the hypothesis that the two pathways contribute in an additive manner to energy generation upon short term glucose starvation, we observed that after 19 h of starvation the double deletion strain showed lower levels of intracellular ATP than each of the single deletion mutants (Figure 6ac) and that survival deficiency increased slightly after 14 days of starvation. We conclude that lipid degradation and autophagy work in parallel to ensure energy maintenance upon glucose starvation.

## Discussion

Cells commonly experience sudden changes in nutrient availability; therefore, strategies to overcome nutrient scarcity are critical to ensure survival in periods of starvation. However, the immediate metabolic response to starvation remains poorly characterized. Here, we aimed to identify the main immediate energy resources of budding yeast upon acute glucose starvation. We showed that respiration is needed for survival and energy maintenance upon glucose starvation, and we examined the metabolic resources that are fed into the respiratory chain. We found that within seconds of glucose starvation, upper glycolysis metabolites decrease drastically. Furthermore, we show that autophagy and β-oxidation are critical energy providers as early as within the first day, and play a central role for survival of yeast during long term starvation.

The role of autophagy in starvation for glucose-grown cells is controversial. A few studies conclude that autophagy is neither activated nor necessary for short-term survival during starvation (12, 32); whereas, other studies suggest the opposite (8, 33). We observed that autophagy is essential for cell survival after glucose starvation within 7 days and that ATP levels depended on autophagy within the first 24h of starvation. While Adachi et al. suggest that autophagy is not induced within a few hours of glucose starvation, they also show that within days, autophagy becomes important for survival of these cells. Our data agrees with the latter, and additionally suggests that energy levels, as measured by ATP, decrease in autophagy-deficient cells already within 19 h.

In the respiratory-deficient *cbp2Δ* mutant, we found sedoheptulose-7-guanine (S7P), guanine, and phosphoenolpyruvate (PEP) to deplete during starvation in direct contrast to wild-type cells. S7P, guanine, and PEP are all potentially connected through gluconeogenesis, pentose phosphate pathway, and nucleotide synthesis. Low PEP concentration would suggest that the *cbp2Δ* mutants have lower gluconeogenic flux (34), which also correlates with the decreased ribose levels observed for rapid timescales (10-60 sec) in the Antimycin A treated starved cells. High concentration of PEP is needed in order to drive the pathway to glucose-6-phosphate and consequently through the pentose phosphate pathway (where S7P resides) eventuating into nucleotides (e.g. guanine). Gluconeogenic synthesis of PEP is primarily catalyzed by PEP carboxykinase. In yeast this reaction is driven by GTP hydrolysis. Given the lower levels of energy cofactors and presumably of energy fluxes in *cbp2Δ* (Figure 1a), the production of PEP by carboxykinase is likely downregulated (35).

An earlier study concluded that μ-lipophagy is triggered by glucose deprivation via the global energy regulator AMPK (8) within several days of starvation. Yeast AMPK activity is known to directly correlate with increased AMP/ATP ratio (36). We had observed changes in the energy charges of the cells (e.g. ATP, ADP, and AMP) (Figure 3e) congruent with the observations of Seo et al. Strikingly, we measured ATP to deplete within seconds while AMP increased in a near equivalent time frame. This activity may suggest that AMPK potentially activates earlier and conveys the intracellular signaling to manage starvation within seconds. While AMPK signaling has been long studied, the kinetics of induction have not been entirely revealed (37). Our data open up the possibility that AMPK signaling may be activated within seconds of starvation.

One of the expected responses from AMPK signaling is an increase in β-oxidation activity. While a direct inhibition of Atg14 dependent μ-lipophagy did not affect the general energy status within the first 19 h of starvation, we found that the ability to perform general β-oxidation is important for ATP maintenance as well as survival in acute glucose starvation, and that lipid intermediates increase within seconds after glucose starvation. Our lipidomics dataset suggests that these immediate changes are not due to large-scale membrane remodeling since we only find that few lipid traces increase upon glucose starvation, in particular polyketides and dolichols (Figure 5d). Polyketides are a structurally very heterogeneous class of compounds of which many have been attributed antimicrobial function (38). The cells might manufacture the polyketides as an antimicrobial measure during starvation. Less microbial competition for remaining nutrients as well as nutrient freeing by eliminating potential competitors could be a rational strategy to ensure survival during starvation.

The other lipid that increased during 30 min of starvation was dolichol, which is required for protein glycosylation in the ER (Figure 5d). Here it functions together with UDP-glucose as a carrier to deliver substrates for glycosylation (39). Interestingly, we also observed the depletion of UDP-hexose (presumably UDP-glucose) in *cbp2Δ* cells within 1-4 h (Figure 1b). In *cbp2Δ* cells all potential intracellular energy sources, including UDP-hexoses, might be drained within hours of starvation due to the lack of efficient energy generation via respiration, while wild type cells might refrain from degrading metabolites needed for rapid re-growth after starvation. The increase of dolichol in wild type cells within 30 min of starvation on the other hand could point to an inhibition of glycosylation upon carbon starvation. Indeed, previous evidence suggests that the dolichol pathway is transcriptionally downregulated in response to glucose starvation (40, 41).

In this manuscript, we sought to better characterize the strategies yeast cells employ as they enter starvation. Our results show that yeast starvation is multifaceted and entails responses at many levels (e.g. metabolites, energy level) and pathways (beta-oxidation, autophagy). We conclude that multiple metabolic responses are needed to ensure sufficient supply of substrates for respiration as the cells maximize their survivability.

## Methods

### Strains and growth

All strains used in this study have a W303 background, precise genotypes are listed in supplementary table 1. Yeast strains were grown in SCD medium (synthetic complete with 2% (w/v) glucose, containing 6.7g/l yeast nitrogen base (Difco) and amino acids according to supplementary table 2) at pH 5.0 (titrated with HCl/KOH) at 30°C rotating. Gene deletions were performed by homologous recombination of a PCR amplified cassette encoding antibiotic resistance, functional amino acid encoding genes, or functional nucleotide encoding genes according to supplementary table 1 (42).

### Acute starvation

Cells were washed 3 times into SC medium (synthetic complete medium without glucose) by repeating the following steps three times: 1. centrifuging for 1 min to pellet the cells, 2. removing supernatant, 3. resuspending the cells in SC medium. Control cells in SCD were treated the same, washing three times into SCD medium.

### ATP measurements

ATP measurements were done according to (8) with minor modifications. Cells were pelleted and resuspended in 750ul 90% acetone. Normalization was done according to OD. The cells were then incubated at 90°C for approx. 10 min to evaporate the acetone, when about 50ul of solution remained. 450ul of buffer (10mM Tris pH 8.0, 1mM EDTA) was added to the solution and ATP was measured with an ATP Determination Kit (Thermo Scientific) on a CLARIOstar microplate reader (BMG).

### Survival assays

Cells were acutely starved and incubated at 30°C rotating. Cells were spotted onto YPD plates with high Adenine, starting at an OD of 0.2, and in a 6-8x dilution series, diluting 5x in every step. The spotting plates were incubated at 30°C for 1-2 days. Percentages of survival compared to WT were assessed by counting CFU (colony forming units) per dilution after x days of starvation, compared with the CFUs counted after the same amount of days in starvation of the wild-type.

### MG132 treatment

*PDR5* was deleted in all strains used for MG132 drug treatments. Cells were treated with 100 μM MG132 (Sigma Fluka Aldrich) for 60mins at 30°C before measuring. For MG132 starvation treatment, cells were pretreated as described with MG132 and then washed 3 times into SC medium containing MG132.

### Fast filtration wash, sampling, and extraction

For each measurement, approximately 1 OD unit of cells (OD600*mL) were captured onto filter paper using a fast filtration set up (21, 22). Immediately, the cells were suffused to flowing pre-treatment media (SCD media) for at least 10 seconds. Upon media switch, the flow of pre-treatment media was stopped, and the post-treatment media was followed for a given amount of time (SC media for 10, 20, 30, and 60 seconds or SCD for 0 and 20 seconds). After the post-treatment media, the cells and filter paper were immediately quenched in 4 mL of extraction solution of 40:40:20% acetonitrile: methanol: water at −20°C. The extraction solution with cells were incubated at −20°C overnight, and the extraction mix was subsequently transferred to 15 ml tubes. The extraction solvent was evaporated at 0.12 mbar to complete dryness in a SpeedVac setup (Christ, Osterode am Harz, Germany), and the samples were dissolved in 100 μl of water and transferred to a 96 well plate for measurement. Samples were stored at −20°C until measurement.

### Metabolomics measurement and annotation

For untargeted analysis (Figures 3a-c), extracts were measured using FIA-QTOF in negative ionization mode and annotated to metabolites as described in antecedent study (20). For targeted analysis (Figures 1b, 2a, 3de), measurement and analysis is described in previous work (24). For all targeted analysis measurements, an internal standard of fully labeled C^13^ extract was used to normalize all data (43).

### Lipidomics extraction

For each sampling, approximately 25 OD units of cells (OD_600_*mL) were captured onto filter paper using a fast filtration set up. The cells with filter paper were immediately quenched in 10 ml of yeast lipid extraction solution (15:15:5:1:0.018 % of ethanol, water, diethyl ether, pyridine, and 4.2N ammonium hydroxide respectively) at −20°C. The extraction solution with cells was incubated at −20°C overnight, and the extraction mix was subsequently transferred to 15 ml tubes. The extraction solvent was evaporated using a SpeedVac setup (Christ, Osterode am Harz, Germany), and the samples were dissolved in 100 μl of 45:45:10 % of isopropanol, methanol, and water respectively and transferred to glass vials with inserts. Samples were stored at −20°C until measurement.

### Lipidomics measurement

Chromatographic separation and analysis by mass spectrometry was done using a 1200 series HPLC system with a Phenomenex Kinetex column (1.7 μl × 100 mm × 2.1 mm) with a SecurityGuard Ultra (Part No: AJ-9000) coupled to an Agilent Technologies 6550 Accurate-Mass Q-Tof. Solvent A: H2O, 10mM formic acid; Solvent B: acetonitrile, 10mM formic acid. 10 μl of extract were injected and the column (C18) was eluted at 1.125 ml/min. Initial conditions were 60% solvent B: 0-2 min, 95% B; 2-4 min, 60% B; 4-5 min at initial conditions. Spectra were collected in negative ionization mode from 50 – 3200mz with high resolution at 4 GHz. Continuous infusion of calibrants (Agilent compounds HP-321, HP-921, HP-1821) ensured exact masses over the whole mass range. We converted the raw data files to the mzML format using msConvert and processed them in R using the XCMS (ver 3.0.2) and CAMERA (ver 1.34.0). M-H and M+FA-H ions were annotated using LIPID MAPS (vers. March, 2017) with a mass tolerance of 0.005 amu (27).

## Acknowledgements

We are grateful to the Sauer and Weis laboratory members for feedback and advice. We would like to thank Elisa Dultz, Stephanie Heinrich, and Ruchika Sachdev for critical reading of the manuscript, Marieke F. Buffing and Brendan Ryback for fast filtration help, and Marieke F. Buffing and Tobias Fuhrer for targeted mass spectrometry help. This work was supported by the Swiss National Science Foundation (SNF 31003A_179275 to K.W.).

## Author contributions

C.W., K.S., U.S., and K.W. conceptualized and organized the project and wrote the manuscript. C.W. and K.S. cultivated the cells. C.W. designed, performed, and analysed all survivability assays and ATP luminescence-based assays, as well as performed the experiments in Figure 1 and 2 except for the mass spectrometry. C.W., K.S., J.T., and U.S. designed the mass spectrometry experiments. K.S., C.W., and P.W. prepared extracts for mass spectrometry measurement (both metabolomics and lipidomics). K.S. and P.W. performed mass spectrometry measurements and analysed the resulting data.

---

**Figure S1.**
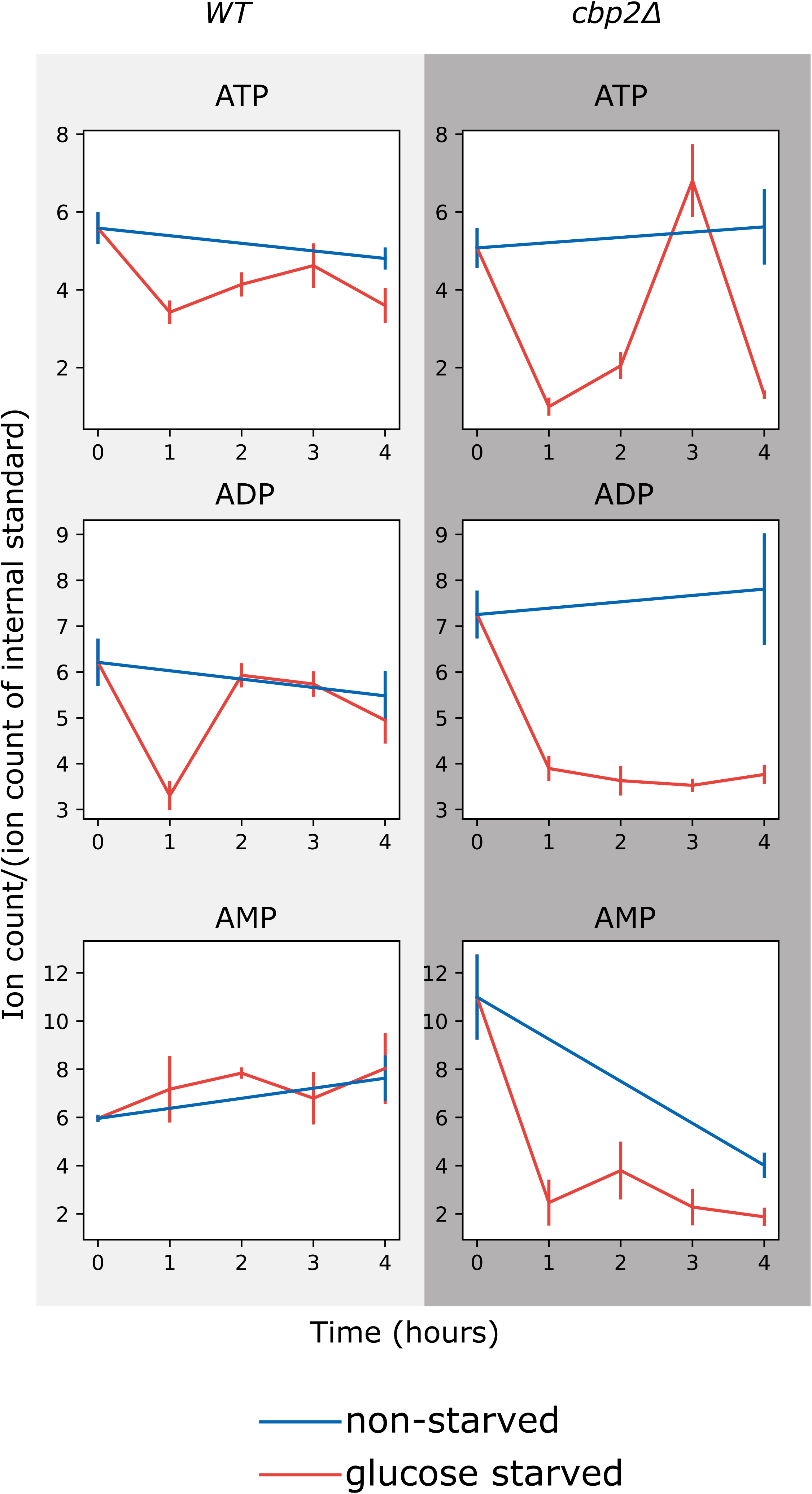
Relative change in energy charges during starvation. Change in ion intensity for ATP, ADP, and AMP after 0, 1, 2, 3, and 4 h of acute glucose starvation (red, glucose starved), compared to non-starved cells (blue, non-starved). Comparison between wild-type (WT) and *cbp2Δ* cells. Average and standard error (error bar) of 2 biological replicates are shown.

**Figure S2.**
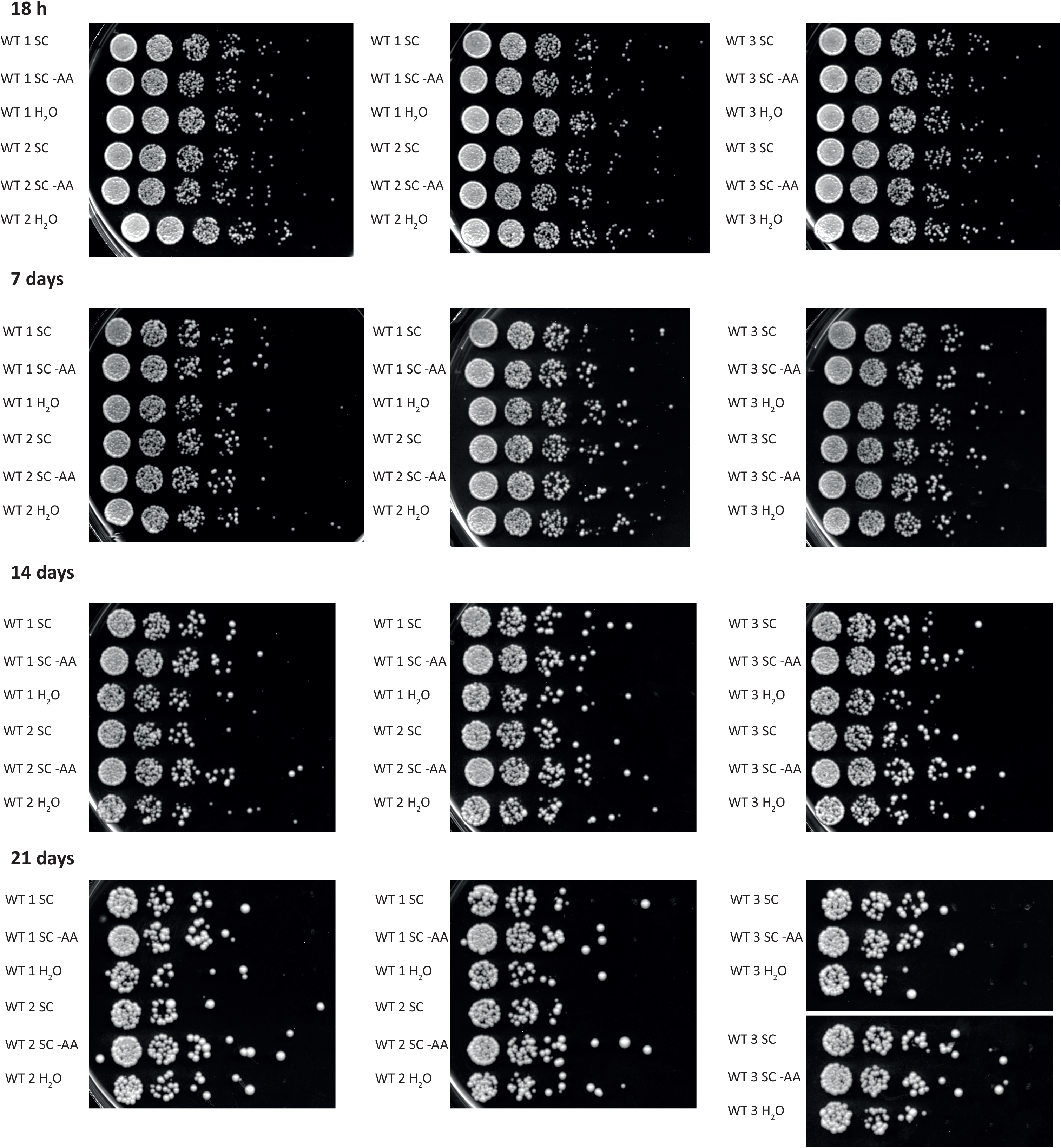
Survival of wild-type yeast cells after 18 h - 21 days of acute glucose starvation (SC), acute glucose and amino acid starvation (SC -AA), and acute starvation in water (H_2_O). 3 biological replicates (WT 1, WT 2, WT 3) and two technical replicates are shown.

**Figure S3.**
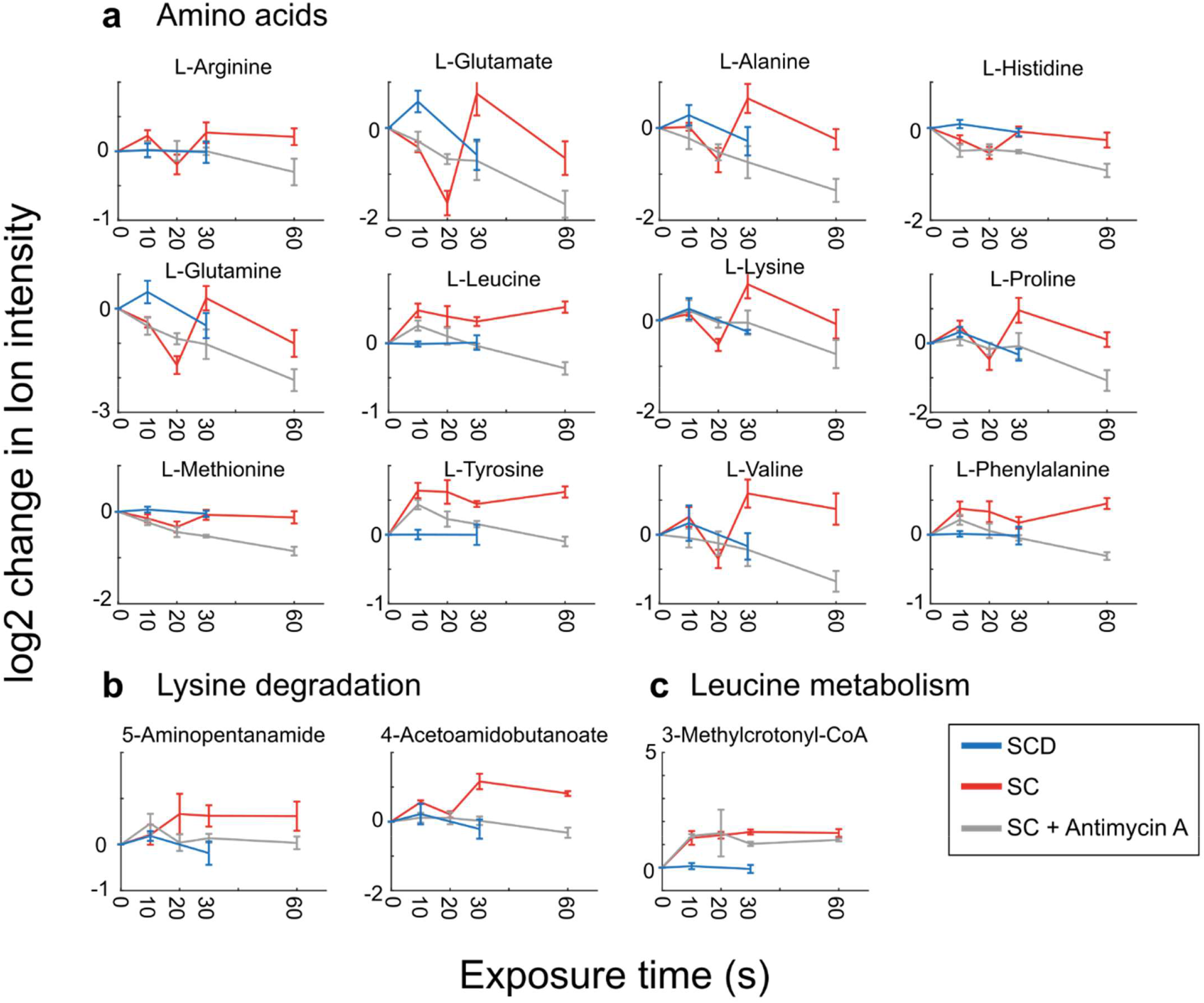
Additional measured ions during rapid starvation. The log2 adjusted values of intensity for specific ions are shown and labeled according to the annotated compound. Measurement time indicates the exposure time for the given media for the cells on the filter (SCD - blue; SC -red; SC with Antimycin A - gray). 6 dots are shown for each timepoint (3 biological replicates and 2 technical measurement replicates). Timepoint 0 s is extrapolated from the average of the SCD condition. Error bars indicate the standard error for three biological replicates.

**Figure S4.**
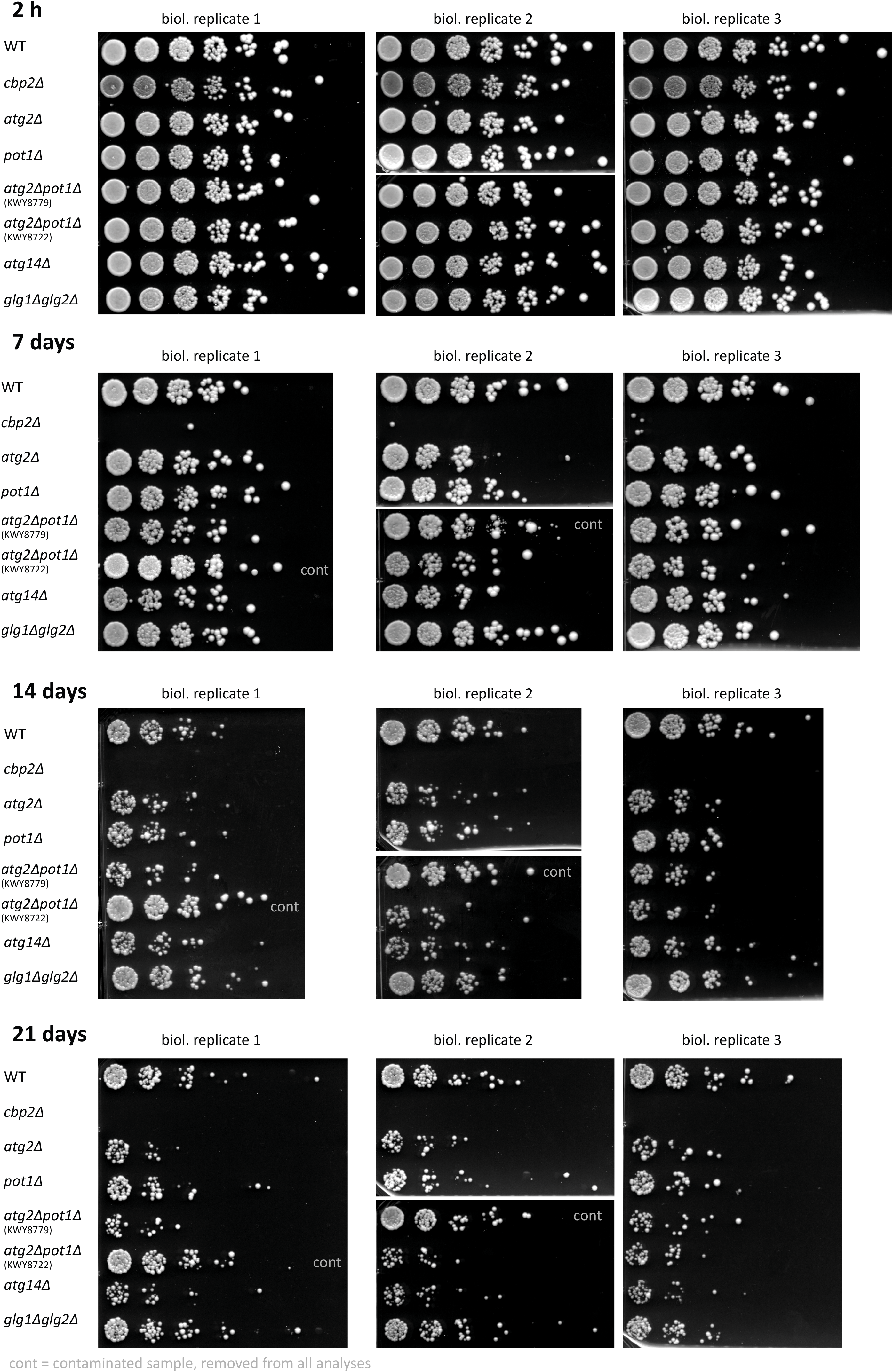
Survival asssays after 2 h – 21 days of acute glucose starvation. 3-4 biological replicates are shown.

## Supplementary Information

**Supplementary Data 1: All metabolomics data**

**Supplementary Table 1:** Strains used in this study.

| Strain | Genotype | Source |
| --- | --- | --- |
| KWY165 | W303 MAT A *his3-11,15 ura3-1 leu2-3 trp1-1 ade2-1* | (9) |
| KWY8358 | W303 MAT A *his3-11,15 ura3-1 leu2-3 trp1-1 ade2-1 cbp2Δ::KanMX6* | This Study |
| KWY7617 | W303 MAT A *his3-11,15 ura3-1 leu2-3 trp1-1 ade2-1 pot1Δ::KanMx6* | This Study |
| KWY8357 | W303 MAT A *his3-11,15 ura3-1 leu2-3 trp1-1 ade2-1 atg2Δ::KanMX6* | This Study |
| KWY8722 | W303 MAT A *his3-11,15 ura3-1 leu2-3 ade2-1 atg2Δ::KanMX6 pot1Δ::TRP* | This Study |
| KWY8779 | W303 MAT A *his3-11,15 ura3-1 leu2-3 ade2-1 pot1Δ::KanMX6 atg2Δ::TRP* | This Study |
| KWY8140 | W303 MATa *his3-11,15 ura3-1 leu2-3 trp1-1 ade2-1 glg1Δ::HygNT1 glg2Δ::KanMX6* | This Study |
| KWY8794, KWY8795, KWY8796 | W303 MAT A *his3-11,15 ura3-1 leu2-3 trp1-1 ade2-1 atg14Δ::KanMX6* | This Study |
| KWY8776, KWY8777, KWY8778 | MAT A *15 ura3-1 leu2-3 trp1-1 ade2-1 Δpdr5::HIS* | This Study |

**Supplementary Table 2:**

|  | mg/l |
| --- | --- |
| Adenine, Arginine, Histidine, Methionine, Tryptophane, Uracil | 20 |
| Lysine, Tyrosine | 30 |
| Phenylalanine | 50 |
| Leucine | 60 |
| Threonine | 200 |

