## Supplementary_Data for "Lipid digestion and autophagy are critical energy providers during acute glucose depletion in *Saccharomyces cerevisiae*": readme.txt

annotation.xls - the annotation file - contains all of the annotated ions that correspond to metabolites. The column ion number corresponds to the Ion number in the data.xlsx file.

data.xlsx - contains the data for the for the raw ion counts of the media. Time point 0 means before the media switch. 
