## Supplementary_Data for "Lipid digestion and autophagy are critical energy providers during acute glucose depletion in *Saccharomyces cerevisiae*": readme.txt

OverFourHoursLC-QQQ.xlsx contains all the measured metabolites using targeted MS methodology. The condition "MM 1" and "MM 2" is the C13 internal standard, for which all data was normalized to before plotting. 

Examples of the categories: 
'WT 1 pre' means wild-type cells, replicate 1, before resuspension into new media. 
'WT 2 SCD 4h' means wild-type cells, replicate 2, measured 4 hours after resuspension in SCD media.
'cbp 2 SC 2h' means cbp2 mutants, replicae 2, measured 2 hours after resuspension into SC media.
