## Supplementary_Data for "Lipid digestion and autophagy are critical energy providers during acute glucose depletion in *Saccharomyces cerevisiae*": readme.txt

matches_em.csv: Contains all possible annotations of the ions from the two experiments
Yeast_starvation_norm.csv: Contains output from data processing with xcms, raw intensities & qnorm intensities.

matches_em.csv file has column with the name "id". The number connects annotation with respective ion in Yeast_starvation_norm.csv, column "X"
