## Supplementary_Data for "Lipid digestion and autophagy are critical energy providers during acute glucose depletion in *Saccharomyces cerevisiae*": readme.txt

annotationWLnChanges.xls - the annotation file - contains all of the annotated ions that correspond to metabolites. The column ion number corresponds to the Ion number in the ph5nodex_ihn.xlsx and ph5nodexdrop.xlsx files.

ph5nodex_ihn.xlsx - contains the data for the switch from SCD to SC for 10, 20, 30, and 60 seconds. The labels indicate the exposure time to SCD and SC. For example, "'ph_5_no_dex 10; 60" means SCD for 10s followed by 60s of SC. "ph_5_no_dex 30;" means SCD for 30s then quench/extraction. 


ph5nodex_ihn.xlsx - contains the data for the switch from SCD to SC + antimycin A for 10, 20, 30, and 60 seconds
