## Supplementary_Data for "Lipid digestion and autophagy are critical energy providers during acute glucose depletion in *Saccharomyces cerevisiae*": readme.txt

LCQQQfastData.xlsx contains all the measured metabolites using targeted MS methodology. The condition "std" is the C13 internal standard, and was used for normalization in the plotting. 
"10s; 60s" means that the cells were exposed to SCD media for 10s, then switched for 60s in SC media and extracted/quenched.
